# Genomic-guided conservation actions to restore the most endangered conifer in the Mediterranean Basin

**DOI:** 10.1101/2023.11.24.568549

**Authors:** José Carlos del Valle, Montserrat Arista, Carmen Benítez-Benítez, Pedro Luis Ortiz, Francisco J. Jiménez-López, Anass Terrab, Francisco Balao

## Abstract

Species with extremely small population sizes are critically endangered due to reduced genetic diversity, increased inbreeding, and the added threat of hybridization. Genomic tools significantly advance conservation by revealing genetic insights into endangered species, notably in monitoring frameworks. Sicilian fir is the most endangered conifer in Europe with only 30 adult trees spread across an 84-hectare area. Using 20,824 SNPs from RAD-seq employing the silver fir genome assembly and a custom 120 SNP-array, we evaluated genetic diversity, mating patterns, and effective population size in adult trees, 118 natural seedlings, and 2,064 nursery seedlings from past conservation actions. We assessed introgression from neighboring non-native fir plantations and established an intra-population assisted gene flow program selecting the most genetically dissimilar individuals and investigating the outcome through simulations. Genomic analysis unveiled significant genetic diversity among adult Sicilian firs, comparable to non-endangered Mediterranean firs with larger populations. However, the genetic diversity of the forthcoming generation declined due to high self-fertilization, leading to marked inbreeding (Fis = 0.38) and an alarmingly low effective population size (*N*e = 6). Nursery seedling monitoring revealed similar selfing rates but significant introgression (∼50%) from non-native firs. Although intra-population assisted gene flow could help to mitigate genetic loss, it may not alleviate the species vulnerability to imminent environmental challenges, perpetuating the risk of an extinction vortex. Hence, investigating the impact of Sicilian fir population decline and selfing on inbreeding depression, along with exploring the potential of hybrids for genetic load alleviation and future adaptation, is crucial for effective conservation strategies. This study stands as a compelling model for guiding conservation strategies in similarly imperiled species characterized by extremely small populations.

## Introduction

Over the past few centuries, many species of animals and plants have become extinct, while many others teeter on the brink of extinction due to habitat loss, fragmentation, and the reduction of population size to inviable levels (Palomares et al. 2012; Benazzo et al. 2017; Humphreys et al. 2019). Species with extremely small populations present a pressing challenge in contemporary biodiversity conservation, referred to as the ’small population paradigm’ (Norris 2004). These species face an increased risk of extinction due to a confluence of factors, including reduced genetic diversity, increased inbreeding, vulnerability to stochastic events, and susceptibility to environmental pressures (Frankham 2015; Ralls et al. 2018). Hybridization presents an additional peril, endangering the species integrity and exacerbating the risks for its already impoverished populations (Balao et al. 2015). To persist at short-term without facing these detrimental effects, it is imperative to maintain an effective population size above 100 individuals (i.e., the minimum viable population sensu Frankham et al. 2014). Consequently, genetic management has become an essential tool to identify strategies for preserving their genetic integrity, mitigating inbreeding depression, and fostering the adaptive potential needed for their survival and long-term persistence in their respective ecosystems (Hogg et al. 2022). Furthermore, genomic tools facilitate the monitoring of current conservation status and the assessment of outcomes from conservation actions (Flanagan et al. 2018; Humble et al. 2023).

Assisted gene flow (AGF) replacing natural gene flow by guided-outcrossing can thus be a powerful conservation tool to preserve biodiversity, particularly for populations that are experiencing genetic erosion (Pregler et al. 2023). Restoring gene flow into small, isolated populations can increase genetic diversity and fitness (i.e., genetic rescue) reducing the risk of inbreeding depression and ultimately population extinction and, at the same time, accelerate the adaptive potential of species to environmental modifications such as climate change (Aitken & Whitlock 2013; Castilla et al. 2019). Despite the benefits and risks of this practice have been extensively discussed (Frankham 2016; Ralls et al. 2020; Grummer et al. 2022), the practical use of the AGF in the tree conservation remains largely unexplored (Aitken et al. 2015; Browne et al. 2019).

Conifers, such as pines and spruces, have specific conservation needs due to their slow growth, long reproductive cycles, low mutation rates, and specific breeding, pollination, and dispersal systems, requiring a meticulous and sustained approach to population restoration (Givnish 1980; Savolainen & Pyhäjärvi 2007; De La Torre et al. 2017)*. Abies nebrodensis* (Lojac.) Mattei (Sicilian fir) is the most threatened conifer in the Mediterranean Basin, and likely one of the most endangered species in the world. It is categorized as “critically endangered” by the IUCN Red List (Thomas 2017), listed in Appendix I of the Bern Convention and considered a priority species in Annexes II and IV of the 92/43 EC Habitats Directive (Code 9220). This species constitutes an example of a highly vulnerable species due to its very low population size, with only 30-adult trees and 118 seedlings (over 5-yr old) growing in a sole population over approximately 84 hectares within the integral reserve of Parco delle Madonie in Sicily (Venturella et al. 1997). Previous studies suggested that reproductive trees maintain high genetic diversity but typically exhibit reduced seed viability (Vicario et al. 1995; Vendramin et al. 1996; Parducci et al. 2001; Scialabba 2019). This low crop viability strongly indicates the existence of high rates of inbreeding. Moreover, fir species show a pronounced tendency for hybridization (Klaehn & Winieski 1962; Balao et al. 2020). The presumed hybridization between the Sicilian fir and the non-native silver and Greek firs (*A. alba* and *A. cephalonica*, respectively) from the surrounding areas of the natural population may adversely impact seed viability and species integrity (Conte & Cristofolini 2003; Scialabba et al. 2005).

Many *in-situ* and *ex-situ* conservation efforts have been implemented over the last two decades, encompassing breeding programs generating thousands of seedlings via natural pollination and the removal of non-native fir species from the vicinity of the population (Venturella et al. 1997; Frascella et al. 2022; Rogatis et al. 2022). However, there is scarce genetic information available on both the natural occurring and the nursery-propagated seedlings, hindering the assessment of a viable long-term conservation plan. To address this limitation, we conducted a genome-wide analysis of Sicilian fir genetic variation and placed our results in the context of genetic diversity observed in other Mediterranean fir species. We then developed an SNP-array for genetic monitoring of the Sicilian firs. We evaluated the genetic diversity and relationships among the 30 adult trees in the natural population. Through simulations, we designed an AGF program that prioritizes crosses using individuals with greater genetic dissimilarity, with the goal of ensuring the short-term survival. We determined the parentage origins of 118 naturally regenerated seedlings, along with 2,064 seedlings cultivated in the forest nursery ’Piano Noce,’ aiming to ascertain outcrossing rates, and occurrences of self-fertilization. Finally, we assessed the degree of introgression with neighboring plantations of non-native firs.

## Materials and methods

### Abies species sampling and RAD-seq libraries

We used restriction site associated DNA sequencing (RAD-Seq) to identify high-quality and information-rich SNPs for genotyping of Sicilian firs and detect hybridization with alien silver and Greek firs. We sampled five individuals from the unique population of Sicilian fir, 15 individuals from three Italian silver fir populations and 15 individuals from three populations of Greek fir (Supplementary Material; Table S1). We extracted DNA and prepared RAD libraries following Balao et al. (2020). Genomic DNA was digested by using high-fidelity SbfI (New England Biolabs) and the resulting fragments were double barcoded. The library was sequenced in a separate lane of an Illumina flowcell HiSeq 2500 at the VBCF NGS Unit (www.vbcf.ac.at/ngs) as 100 bp single-end (SE) reads.

### SNP identification and population genomics

Quality filtering and demultiplexing of the RAD-seq library were performed with deML (Renaud et al. 2015) and STACKS ver. 2.53 (Catchen et al. 2011, 2013, Rochette et al., 2019). Following this, we aligned each raw read fastq file to the silver fir genome (Mosca et al 2019), using Bowtie2 (Langmead & Salzberg 201) with --sensitive and --no-unal settings. Single-best alignments were then sorted and converted from SAM to BAM format using Samtools v1.10 (Li et al., 2009). To assemble RAD loci, we used the ref_map.pl pipeline implemented through Stacks v.2.53. Then, we utilized the populations script to export SNP data into various formats for subsequent analyses. We investigated the nucleotide diversity (π), heterozygosity (*H_s_*) and inbreeding (*F_IS_*) of the Sicilian fir and compared to silver and Greek firs populations.

### OpenArray development for genotyping and hybrid detection

We customized a panel of 120 SNPs by PCR-based OpenArray Technology (Thermo Fisher Scientific, USA) for genotyping the Sicilian firsamples. Firstly, we selected 100 highly informative SNPs to assess the genetic structure of the natural population as well as the origin of the seedling from the forest nursery ‘Piano Noce’. These unlinked high-quality SNPs with a maximum information for genotyping and paternity exclusion were selected using a combination of Stacks, Plink2 v. 2.00a3.7 (Chang et al., 2015), PopLDdecay 3.40 (Zhang et al., 2019) based on the following criteria: ‘Biallelic’, ‘No other SNPs within 4 kb’, ‘*r*^2^ < 0.01’, ‘Minor allele frequency [MAF] > 0.05’, ‘Minimum 1 individual heterozygote’, ‘Minimum 1 individual homozygote for alternative alleles’, ‘100 bp flanking sequence available on both sides of the variable position’. To minimize the probability of SNPs being affected by natural selection, we kept the SNPs in intergenic regions (avoiding 1kb downstream and 2 kb upstream regions of genes) using SnpEff and SnpSift v4.3.1t (Cingolani et al., 2012). Secondly, we selected 20 high-quality SNPs (unlinked markers with high MAF) to detect hybridization between the Sicilian fir and the other firs inhabiting the Madonie Natural Park. These selected makers followed the same selection criteria previously described and showed the highest species-specific diagnostic power following the highest loadings in the first two axes of a Discriminant Analysis of Principal Components (DAPC) performed with the *adegenet* package v. 2.1.10 (Jombart, 2008) in R v.4.0.3 (R Core Team 2020). A Principal Component Analysis (PCA) using the R package *dartR* 2.7.2 (Mijangos et al., 2022) was used to corroborate the discriminant power of the 20 selected SNPs. We customized a panel of 120 SNPs by PCR-based OpenArray Technology (Thermo Fisher Scientific, USA) for genotyping Sicilian fir samples. Firstly, we selected 100 highly informative SNPs to assess the genetic structure of the natural population as well as the origin of the seedling from the forest nursery ‘Piano Noce’. These unlinked high-quality SNPs with a maximum information for genotyping and paternity exclusion were selected using a combination of Stacks, Plink2 v. 2.00a3.7 (Chang et al. 2015), PopLDdecay 3.40 (Zhang et al. 2019) based on the following criteria: ‘Biallelic’, ‘No other SNPs within 4 kb’, ‘*r*^2^ < 0.01’, ‘Minor allele frequency [MAF] > 0.05’, ‘Minimum 1 individual heterozygote’, ‘Minimum 1 individual homozygote for alternative alleles’, ‘100 bp flanking sequence available on both sides of the variable position’. To decrease the probability of selecting SNPs under selection, we kept the SNPs in intergenic regions (avoiding 1kb downstream and 2 kb upstream regions of genes) using SnpEff and SnpSift v4.3.1t (Cingolani et al. 2012). Secondly, we selected 20 high-quality SNPs (unlinked markers with high MAF) to detect hybridization between the Sicilian fir and the other firs occurring at the Madonie Regional Natural Park. These selected makers followed the same selection criteria previously described and showed the highest species-specific diagnostic power following the highest loadings in the first two axes of a Discriminant Analysis of Principal Components (DAPC) performed with the *adegenet* package v. 2.1.10 (Jombart & Ahmed 2011) in R v.4.0.3 (R Core Team 2020). A Principal Component Analysis (PCA) using the R package *dartR* 2.7.2 (Gruber et al. 2018) was used to corroborate the discriminant power of the 20 selected SNPs.

### SNP-Array validation and Genotyping of plants from the natural population

To investigate the genetic diversity and relatedness of Sicilian fir individuals, we first collected leaf material for genetic analysis from the 30 adult trees and the 118 seedlings inhabiting the population. DNA was isolated using the NucleoMag Plant kit (Macherey-Nagel, Germany) according to the manufacturer’s protocol. Before starting the genotyping of samples, we initially validated the specially developed 120-SNPs arrays for this species. To do so, we genotyped 24 samples (12 in duplicate) and we assessed replicability and genotype accuracy, as well as we calculated the genotyping error rate and allele drop-out. Then, we carried out a PCA to verify that replicates clustered together.

We used the 120 SNPs panel to genotype the individuals in the natural population. We explored the population genetic structuring of the natural population using Structure v. 2.3.4 (Pritchard et al. 2000). We estimated the number of genetic clusters (*K*) by assigning individuals in undefined mixture clusters under a Bayesian framework. We conducted 10 independent runs of 1,000,000 iterations each one, with a burn-in period of 100,000 for each value of *K* from 1 to 5. Then, the best *K* that fit the data was calculated using the Δk method (Evanno et al. 2005) in Structure Harvester (Earl & VonHoldt 2011). Finally, the effective population size (*N_e_*) and inbreeding level (*F_IS_*_)_ were estimated using the software Colony v.2.0.6.6 (Jones & Wang 2010).

We inferred the pedigree relationships in the natural population of *A. nebrodensis*, using Colony2 with the full-likelihood approach. Our analysis incorporated prior information on parental (all reproductive adults) and maternal genotypes (the closest adult to the seedling). We assumed monoecy, diploidy, both male and female polygamy, and potential inbreeding. For the full likelihood calculation, we employed very high precision setting, very long run length, sibship scaling, updated allele frequencies, and weak sibship priors. The genotyping error rate (estimated from replicates) was set to 0.01 for all loci. We performed three runs with different random number seeds to check the reliability of the results. Additionally, the seedlings with a putative hybrid origin (i.e., with an uncertain genetic origin) were examined with a PCA using the 20 SNPs designed for the discrimination of fir species.

For the adult trees, we used the *adegenet* package to estimate the genetic diversity at the individual level by calculating the following coefficients: *HsExp*, standardized heterozygosity based on the mean expected heterozygosity; *HL*, homozygosity by loci; and *INBR*, inbreeding coefficient. Additionally, we used Colony2 to estimate the effective population size (*N_e_*) and inbreeding level (*F_IS_*_)_. To assess whether genetic information could be used in management strategies to mitigate the alleged low population diversity/inbreeding, we studied the population genetic structure and relatedness of the 30-adult-trees to design a genome-informed assisted gene flow (AGF) program. To do so, we used the *adegenet* package to perform a DAPC using a *K*-mean clustering (*K* = 1-30, the total number of adult trees). Additionally, we used the *related* package (Pew et al. 2015) to calculate the Ritland estimator (RIT) of pairwise co-ancestry (Ritland 1996) to infer the genetic relatedness of all adult trees. Finally, considering the absence of other Sicilian fir populations for outcrossing, we selected the 12 pairs of adults with the lowest RIT values (i.e., those individuals genetically more dissimilar) to simulate an artificial population in the AGF conservation program. We used the function *hybridize* from the *adegenet* package to generate 10 progenies from each of the 12 proposed crosses. Then, we compared the genetic diversity, inbreeding and effective size of adults, natural seedlings, the simulated AGF population (n = 120) and the AGF next generation population, which is composed by the 120 seedlings from the AGF population, the 118 natural seedlings and 19 seedlings from the nursey originate by outcrossing (see Results section).

### Pedigree analysis of seedlings from the forest nursery

We investigated the genetic diversity and pedigree from 2,064 seedlings at the forest nursery from eight maternal batches from the nursery. To assess a feasible parentage with any of the 30-adult trees and detect putative hybrids with other alien firs, we genotyped the samples with the 120 SNP panel and then we conducted paternity tests and determined the rate of outcrossing, inbreeding and self-fertilization using Colony2, as described above.

## Results

### Genomic diversity of the Sicilian fir and relatives

The RAD-seq of the 35 individuals from populations of Sicilian, silver and Greek firs produced a final STACKS catalogue a total of 365,582 RAD loci (composed of 34,325,726 genomic sites) with an average coverage per sample (± s.d.) of 51.4 ± 38.5 reads per locus. After filtering for polymorphic RAD loci with MAF > 0.05, we obtained a final data set with 20,824 SNPs. The raw data was deposited in the NCBI Short Reads Archive (BioProject ID PRJNA563575).

Estimates of genetic diversity in the populations were notably consistent across the three fir species (Figure 1). Although the Sicilian fir exhibited the lowest values for π and *H*_s_, these differences were not statistically significant when compared to the populations of the other two firs (Kruskal-Wallis tests with *p* > 0.05). The Sicilian fir population showed similar *F_IS_* to those of the silver (0.07 ± 0.014) and Greek (0.06 ± 0.007) firs. Furthermore, it is noteworthy that the Sicilian fir displayed the highest number of private alleles, with 1029, surpassing the populations of the other species which ranged from 483 to 728 (Figure 1). The pairwise *F’_ST_* values varied between 0.07 (silver fir pop1-silver fir pop2) and 0.26 (Sicilian fir – Greek fir pop 2). Finally, the Sicilian fir showed the lowest *F’_ST_* pairwise with the silver fir pop1 (0.14; Table S2).

**Figure 1.**
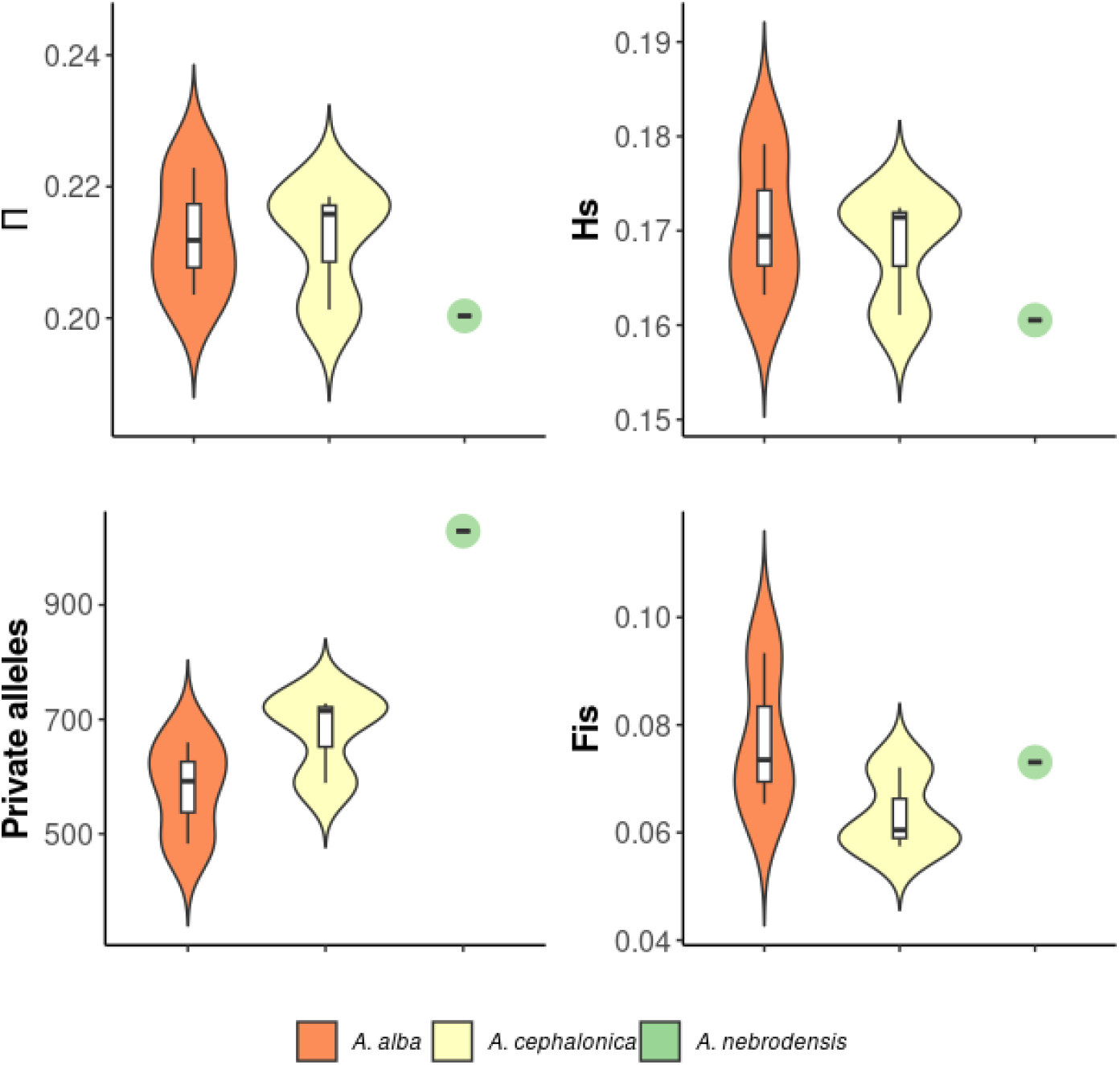
Violin plots display genetic diversity estimators based in 20,824 SNPs for silver (orange), Greek (light yellow), and Sicilian (green) firs. Metrics include nucleotide diversity (π), heterozygosity (*H*s), private alleles, and inbreeding (*F*_IS_).

### Development of SNP-Array for conservation of Sicilian fir

We selected 120 high-quality SNPs from the 3,536 SNPs called with MAF > 0.05 present in the five individuals of the Sicilian fir, remaining only 459 SNPs after the quality filtering. Functional annotations showed that 351 SNPs were in intergenic regions, and 268 SNPs remained after excluding promotors genes regions (46 downstream and 59 upstream). The 20 SNPs selected to discriminate putative hybrids maximized the separation of the three fir species (the first two components of the PCA accounted for 42.5% and 20.8% of the total variation, respectively; Figure S1). The validation of the designed 120 SNP-array revealed a replicability higher than 99% and the PCA successfully separated samples and clustered duplicates together. Call rates (i.e., proportion of samples that were assigned a genotype call compared to the total number of samples) of the analyzed samples ranged from 64.6% to 77.1%. High genotyping failures (> 50%) were detected at 15 loci and were discarded for downstream analyses.

### Gene diversity and population structure in the natural population

The STRUCTURE analysis showed that the most likely number of genetic groupings was *K* = 2 (Figure 2) with no clear clustering pattern. The first cluster comprised three adult trees and 41 out of the 42 seedlings related to one of these trees. The second cluster included the remaining adult trees and seedlings from the population. The effective population size was very low (*N_e_*= 6; 3–21; lower and upper confidence interval at 95%) and the level of inbreeding was moderate (*F_IS_*= 0.373).

**Figure 2.**
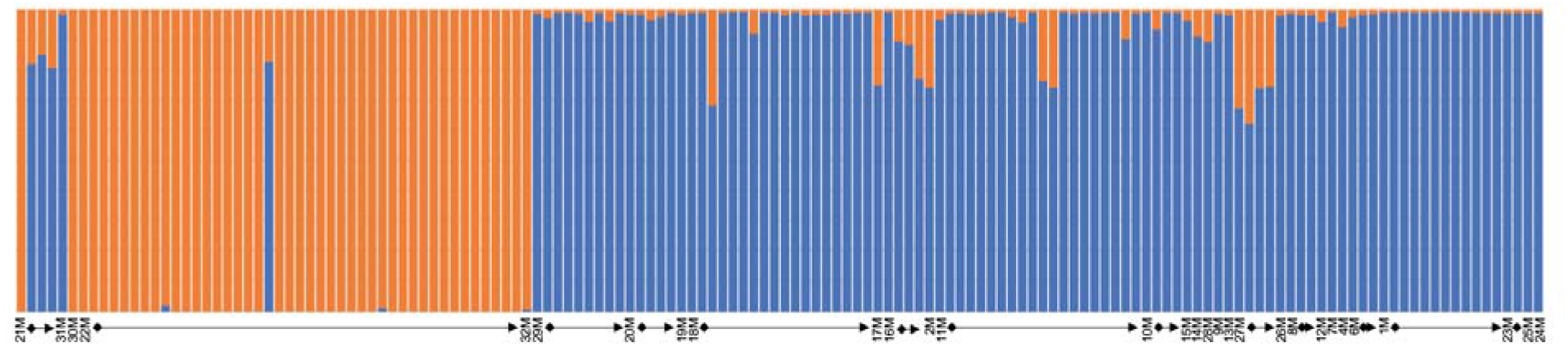
Individual ancestry at *K*= 2, the most supported numbers of clusters in Structure analysis, for the 30 adult trees and the 118 seedlings in the Sicilian fir population. Adults are sorted by proximity and arrows show seedlings origin from mother plants.

The results of the paternity assignments disclosed the potential hybrid origin in nine out of 118 seedlings from the natural population. The genetic origin of three of them could not be assigned to any of the 30-adult trees occurring in the population, and for the remaining six seedlings only one parental was assigned. 109 seedlings (92.4%) were identified as purebred Sicilian firs. Of those, 103 seedlings originated by self-fertilization and only six of them resulted from outcrossing (only five out of the 30 adult-trees were involved in outcrossing). We analyzed in the PCA the nine seedlings with a putative hybrid origin, finding that only one of them was genetically closer to the cloud point of the silver fir and, therefore, it was confirmed as a hybrid (Figure 3).

**Figure 3.**
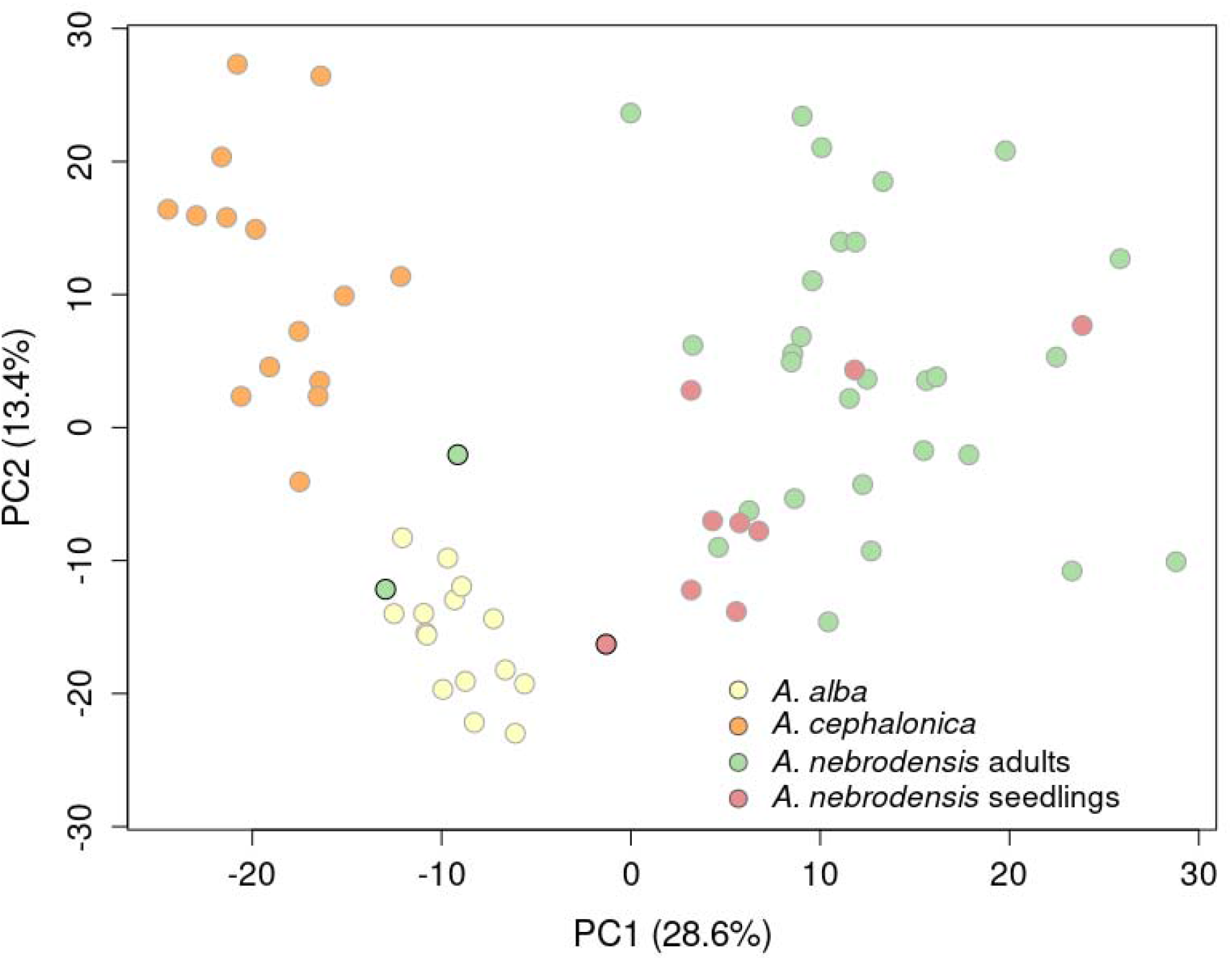
PCA plot displaying nine seedlings of uncertain origin from the natural Sicilian fir population and other alien firs species. Colored dots represent Sicilian (green), silver (light yellow), and Greek (orange) firs, and seedlings with ambiguous origins (red). Darker colors highlight two adult trees and one seedling showing introgression.

For adult trees, the estimations of parental similarity showed mild genetic diversity at the individual level. The mean value of the standardized observed heterozygosity (*HsExp*) was 0.74 ± 0.19, varying from 0.32 to 1.13. The individual homozygosity by loci (*HL*) ranged from 0.50 to 0.86, with an average value of 0.67 ± 0.09. Finally, the inbreeding coefficient (*INBR*) showed values that ranged from -0.20 to 0.79, with an average value of 0.21 ± 0.23. DAPC analysis conclusively found three genetic clusters with almost lacking genetic admixture that were comprised of 12, 11 and seven individuals, respectively (Figure S2A). Co-ancestry analysis using *RIT* pairwise estimations revealed overt differences of the genetic relatedness of the 30-adult trees, ranging from -0.3837 to 1.0587 (a matrix of pairwise co-ancestry among all adult trees is depicted in Figure S2B). Finally, the Mantel test showed absence of any spatial correlation between geographical distance and genetic diversity from adult trees (*r* = -0.06; *p* > 0.05).

### Gene diversity and population structure in seedlings from the forest nursery

We were able to genetically characterize 1,776 of the 2,064 selected seedlings from the nursery. Paternity tests confidently assigned 1,525 seedlings (85.8%) to one or more parents of the Sicilian fir. We identified 897 purebred seedlings from this species with a very high autogamy rate (97.9%), only 19 of them being the result of outcrossing. Additionally, we identified 879 seedlings as putative hybrids. Finally, the effective population size of the seedlings in the nursery was very low (*N_e_*= 12; 6–26; lower and upper confidence interval at 95%) and its level of inbreeding was moderately high (*F_IS_*= 0.354).

### Simulating the effects of the Assisted Gene Flow

RIT values of 12 pairs of adult trees selected to simulate an artificial AGF population ranged from –0.3837 to –0.2843. The simulated AGF population from the selected crosses showed higher *H_o_* (Kruskal-Wallis’s test, *χ^2^* = 91.882, *df* = 3, *p*□<□0.001) and lower inbreeding coefficient (*χ^2^* = 179.82, *df* = 3, *p*□<□0.001) than the natural population (i.e., adults and seedlings) and the next generation population (including natural seedlings, seedlings from the simulated AGF event, and the 19 outbreed seedlings from the nursery). This latter population maintained similar values than current adults in the population whereas natural seedlings showed significatively the lowest *H_o_* and the highest *F_IS_* (Figure 4). The nucleotide diversity was similar for all the groupings (*χ^2^* = 1.984, *df* = 3, *p*□>□0.05). Natural seedlings showed the lowest *N_e_* (95% IC 2.6-2.9) and the adults the highest one (18-25.2). AGP seedlings and the Next-Gen population displayed intermediate *N_e_* values (Figure 4).

**Figure 4.**
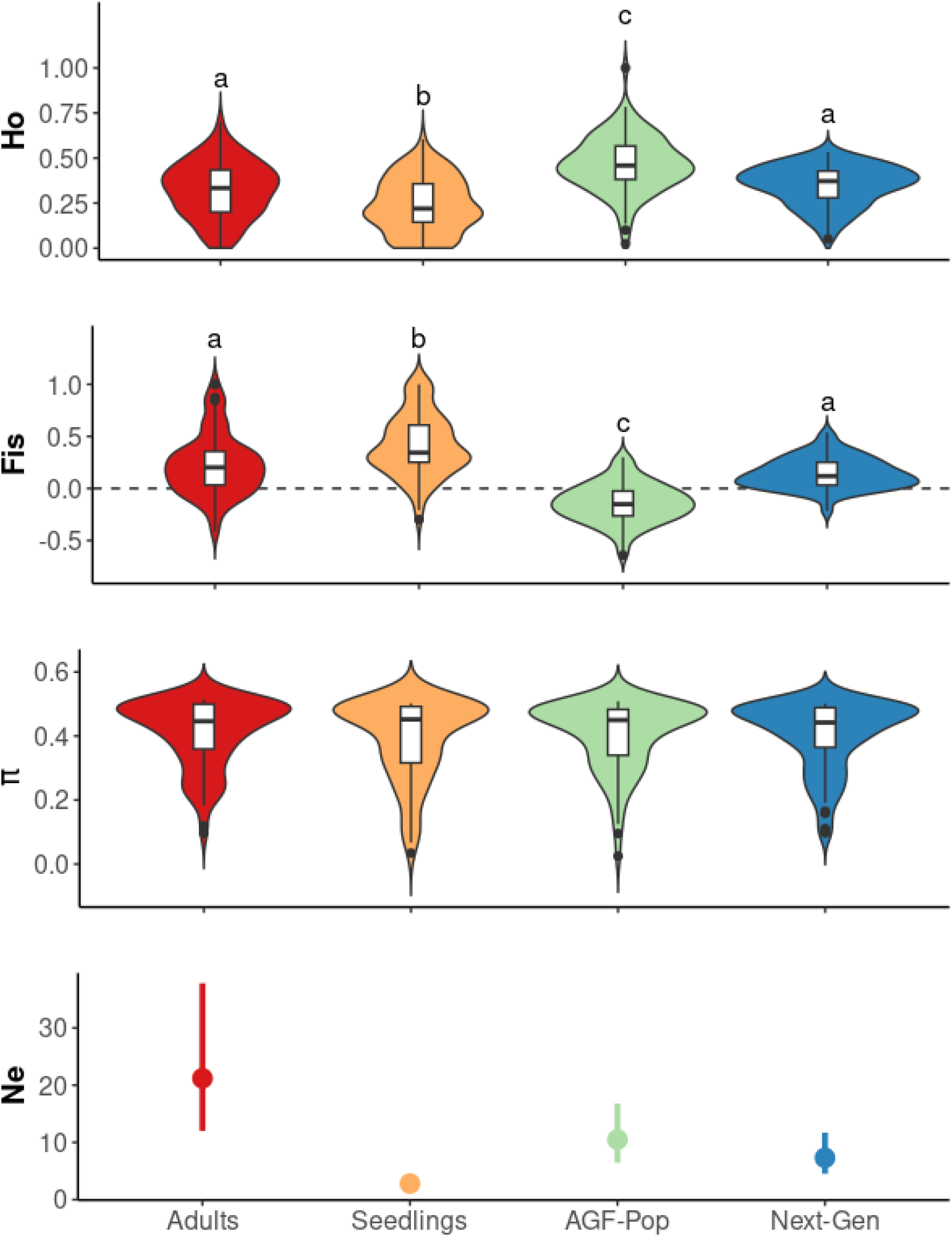
Genetic population parameters based in 120 SNP-array for the adult Sicilian firs (red) and seedlings (orange) from the natural population, the simulated AGF population (green), and the AGF-next-generation population (blue). Violin plots display the distribution of homozygosity (*H_o_*), inbreeding (*F_IS_*), nucleotide diversity (π), and effective size population (*N_e_*).

## Discussion

### Differential genetic diversity across age cohorts of the Sicilian fir

The genome-wide levels of genetic diversity among adult individuals in the Sicilian fir population were comparable to those of the silver and Greek firs populations along their native ranges. The significant genetic variation observed in adult Sicilian fir samples, highlighted in prior studies (Vicario et al. 1995; Ducci et al. 1999; Conte et al. 2004), correlated with the recent demographic dynamics of the Sicilian fir, as evidenced by the increased private alleles, slower heterozygosity response, and stable inbreeding coefficients (Llorens et al. 2018). Palaeobotanical and humanistic studies revealed the massive presence of firs along Sicily from Paleogene to Holocene (Tinner et al. 2016), along with a significant demographic decline since the Middle Ages (Pasta et al. 2020). Anthropogenic pressures nearly decimated the Sicilian fir forests, driving the species to the verge of extinction, with only a few surviving individuals comprising the current population (Venturella et al. 1997). The extended lifespan of firs (> 150 years) and adaptive selection against homozygous individuals could counteract the effects of demographic decline on genetic diversity (Yao et al. 2007; Fraser et al. 2014; Su et al. 2018).

Despite the high genetic diversity observed in adult trees, genetic monitoring of seedlings in the natural population using a developed SNP-array revealed a concerning scenario for species conservation. The seedlings exhibited a substantial decrease in heterozygosity and an increased inbreeding due to exceptionally high selfing rates (> 94%). Only six outcrossed seedlings were found, which were traced back to just five parental genotypes. This contrasts sharply with the typically high levels of outcrossing observed in Mediterranean firs and other gymnosperms (Restoux et al. 2008), and it is likely influenced by several factors such as the sparsely tree dispersion, with an average distance of 621.7 meters between them, and a limited pollen dispersal as occurs in other endangered Mediterranean firs (Arista & Talavera 1994; Sánchez-Robles et al. 2014). The high selfing rate and subsequent decline in genetic diversity across young cohorts of the Sicilian fir likely account for the previously reported reduced seed and seedling viability (Conte et al. 2004; Scialabba 2019; Mirabile et al. 2023). This reduced genetic diversity may also increase the species’ vulnerability to emerging environmental challenges, particularly those associated to climate change (Frankham 2015; Ralls et al. 2020).

### Remnants of recent hybridization in Sicilian fir population and implications in the conservation

It is remarkable that, despite a high level of selfing, at least one seedling in the Sicilian fir population likely derived from hybridization with an alien firs species. Previous conservation efforts identified more than 5,000 such firs in reforested areas located within a few hundred meters of the Sicilian fir population (Ducci 2014). While many of these non-native firs were removed, some still persist, presenting a real risk of genetic exchange, particularly the silver fir, the ancestral species that gave rise to the Sicilian fir (Balao et al. 2020). Hybridization is a widespread phenomenon present in evolutionary history of Mediterranean firs (Krajmerová et al. 2016; Balao et al. 2020). While hybridization can potentially harm endangered species (Balao et al. 2015), understanding the consequences of hybridization requires detailed study (Arnold 2015; Gompert & Buerkle 2016). Notably, two adult Sicilian firs exhibited indications of coancestry with silver firs in the PCA, suggesting historical introgression of anthropical origin due to plantations from at least one century as occurred in other tree species (Vanden Broeck et al. 2005; Meirmans et al. 2014; Scotti-Saintagne et al. 2023). Considering that such introgressed trees could potentially enhance the adaptability towards prevailing ecological stress (Kormutak et al. 2013; Stejskal et al. 2016), it is cautious to prioritize their conservation as part of the Sicilian fir recovery program (Jackiw et al. 2015; VonHoldt et al. 2022; Brauer et al. 2023).

### Monitoring ex-situ regenerated seedlings with SNP-array confirms the prevalence of hybridization and selfing

Similarly, SNP-array analyses of nursery-grown seedlings revealed a 50% prevalence of hybridization, contrasting starkly with the minimal 1-in-118 ratio in natural seedlings. This raises questions about the viability of hybrids in the wild (Campbell & Waser 2001). Clearly, this prevalence of hybrids in the nursery may compromise the implementation of reforestation actions. However, it seems worthwhile to delve deeper into investigating the fitness and adaptability of these hybrids in their natural habitat, considering various environmental factors.

Additionally, approximately 98% of the purebred Sicilian fir seedlings in the forest nursery originated from self-pollination, mirroring the selfing rates observed in the wild. While the effective population size (*N_e_*) in the nursery population, established from cones collected from adult trees, slightly surpassed that of the wild population, this discrepancy likely stemmed from the inclusion of hybrid seedlings. However, the strikingly low ratio of effective population size to census size (*N_e_*/*N*=0.0058) highlights an alarming genetic bottleneck. This compromises the genetic robustness of the forest restoration stock, akin to endangered conifers such as the Boise araucaria or the Yellow Box (Kettle et al. 2008; Broadhurst 2013).

### Implementing within-population AGF as the last chance to avoid extinction

In a similar vein, the Sicilian fir population exhibited a notably low effective population size (*N_e_*=3-21), falling significantly below the recommended threshold (> 100), which poses considerable challenges for its short-term conservation efforts (Frankham et al. 2014). The high selfing rate, coupled with the low mutation rate in firs (De La Torre et al. 2017), indicates that relying solely on spontaneous mutations will not sufficiently enhance the population genetic diversity in the short- or medium-term, emphasizing the urgent need for proactive genetic management strategies (Ralls et al. 2018). Consequently, genetic rescue, through upcoming outcrossing events with genetically diverse individuals, stands as the final opportunity to avert extinction by mitigating inbreeding and reducing levels of homozygosity (Ingvarsson 2001; Fady et al. 2020). Given the single remaining wild Sicilian fir population, we propose a within-population AGF approach, selecting crosses among trees with more distant co-ancestry in the population. This method appears to be effective in enhancing genetic diversity and *N*e, reducing inbreeding, and simultaneously mitigating potential negative effects, such as outbreeding and the introduction of maladaptive alleles associated with AGF (Aitken & Whitlock 2013; Grummer et al. 2022). However, the gain in genetic diversity through AGF restoration appears insufficient to recover past diversity, and the long-term effectiveness of this conservation action to avoid the extinction vortex remains uncertain. Hence, there is a necessity to investigate the impact of Sicilian fir population decline and selfing on inbreeding depression (Pečnerová et al. 2023), along with exploring the potential of hybrids to alleviate genetic load and facilitate future adaptation (vonHoldt et al. 2018).

### Conclusions

Genomic data revealed that adult Sicilian firs harbor high genetic diversity despite their small population size, comparable even to that observed in other Mediterranean firs spread across large populations. Despite this high genetic richness found in adult trees, the genetic variability of the population will dramatically decrease in the upcoming generations, even if human-mediated actions are taken to increase, or at least maintain, the genetic diversity of the population. Intra-population assisted gene flow can contribute to preventing the loss of genetic variation in the population; however, this does not seem to be sufficient to break free from the extinction spiral because of the predictable species’ vulnerability in upcoming generations to emerging environmental challenges, such as the climate change. Interspecific crosses with other intersectional fir species, namely the silver fir, could represent a potential alternative strategy for the future conservation of the Sicilian fir that merits exploration. Our results emphasize the relevance of genomic-guided conservation actions, which can assist in identifying suitable individuals for reforestation and provide a solid foundation for conservation management. We expect that this approach could serve as a model to be replicated in the conservation strategies for other endangered firs, or even for other conifers, around the Mediterranean region.

## Acknowledgments

The authors acknowledge the Ente Parco delle Madonie and the forest nursery of Piano Noce (Polizzi Generosa) for hosting the project area and maintaining *A. nebrodensis* seedlings. We also thank University of Seville Biology and Herbarium General Research Services (CITIUS) for providing the facilities. This research was funded by the European Union through the LIFE4FIR project, grant number LIFE18/NAT/IT/000164.

## Credit authorship contribution statement

**JC del Valle**: Methodology, Investigation, Writing – original draft. **M Arista**: Conceptualization, Methodology, Investigation, Funding acquisition; **C Benítez-Benítez**: Methodology, Investigation; **PL Ortiz**: Conceptualization, Methodology, Investigation; **FJ Jiménez-López**: Methodology, Investigation; **A Terrab**: Conceptualization, Methodology, Investigation; **Francisco Balao**: Conceptualization, Methodology, Investigation, Writing – original draft. All authors have read and agreed to the published version of the manuscript.

---

**Table 1.** Results of the genetic diversity estimations calculated for the 30 adult trees of the Sicilian fir. *HsExp* (standardized heterozygosity based on the mean expected heterozygosity), *HL* (homozygosity by loci), and *INBR* (inbreeding coefficient).

| Adult tree | *HsExp* | *HL* | *INBR* |
| --- | --- | --- | --- |
| 01M | 0.60 | 0.74 | 0.25 |
| 02M | 0.76 | 0.65 | 0.15 |
| 04M | 0.72 | 0.67 | 0.25 |
| 06M | 0.69 | 0.68 | 0.20 |
| 07M | 0.69 | 0.69 | 0.20 |
| 08M | 0.79 | 0.64 | 0.57 |
| 09M | 0.76 | 0.66 | 0.12 |
| 10M | 0.90 | 0.61 | 0.17 |
| 11M | 0.83 | 0.61 | 0.11 |
| 12M | 0.93 | 0.57 | -0.01 |
| 13M | 1.13 | 0.50 | -0.20 |
| 14M | 0.93 | 0.59 | 0.02 |
| 15M | 0.74 | 0.67 | 0.08 |
| 16M | 0.37 | 0.83 | 0.40 |
| 17M | 0.83 | 0.63 | 0.10 |
| 18M | 0.90 | 0.61 | 0.12 |
| 19M | 0.42 | 0.83 | 0.79 |
| 20M | 0.81 | 0.63 | 0.06 |
| 21M | 0.58 | 0.75 | 0.55 |
| 22M | 0.97 | 0.57 | -0.02 |
| 23M | 0.79 | 0.63 | 0.07 |
| 24M | 0.74 | 0.66 | 0.13 |
| 25M | 0.74 | 0.66 | 0.18 |
| 26M | 0.93 | 0.58 | 0.04 |
| 27M | 0.60 | 0.73 | 0.31 |
| 28M | 0.65 | 0.71 | 0.17 |
| 29M | 0.97 | 0.57 | -0.01 |
| 30M | 0.49 | 0.78 | 0.61 |
| 31M | 0.32 | 0.86 | 0.65 |
| 32M | 0.58 | 0.74 | 0.36 |

