## Supplemental Material for "Genomic-guided conservation actions to restore the most endangered conifer in the Mediterranean Basin"

*Departamento de Biología Vegetal y Ecología, Universidad de Sevilla, Sevilla, Spain*

The co-first authors are denoted by an asterisk.

**Table S1.** Population details of the samples included in the RAD-seq study.

| Species | Code | Locality | Country | Altitude (m) | Latitude | Longitude |
|---|---|---|---|---|---|---|
| *A. alba* | ALB_1 | Piano dábete | Italy | 991 | 38.26388 | 16.02539 |
| | ALB_2 | Laurenzana | Italy | 1192 | 40.41872 | 15.97601 |
| | ALB_3 | Selva di Castiglione | Italy | 1005 | 41.73423 | 14.31645 |
| *A. cephalonica* | CEP_1 | Kalavryta | Greece | 633 | 38.07316 | 22.12514 |
| | CEP_2 | Oros Ainos Kefalonia | Greece | 891 | 38.15680 | 20.66164 |
| | CEP_3 | Oros Pateras | Greece | 747 | 38.14394 | 23.31128 |
| *A. nebrodensis* | NEB_1 | Parco delle Madonie | Italy | 1659 | 37.87568 | 14.04049 |

**Table S2.** Pairwise $F_{ST}$' values among Sicilian, Greek, and silver fir populations. Population names as indicated in Table S1.

| | NEB_1 | CEP_1 | CEP_2 | CEP_3 | ALB_1 | ALB_2 | ALB_3 |
|---|---|---|---|---|---|---|---|
| **NEB_1** | 0.0000 | 0.2293 | 0.2577 | 0.2409 | 0.1428 | 0.1635 | 0.1930 |
| **CEP_1** | - | 0.0000 | 0.1056 | 0.1024 | 0.1659 | 0.1737 | 0.1636 |
| **CEP_2** | - | - | 0.0000 | 0.1407 | 0.1933 | 0.2195 | 0.2238 |
| **CEP_3** | - | - | - | 0.0000 | 0.1902 | 0.2003 | 0.1864 |
| **ALB_1** | - | - | - | - | 0.0000 | 0.0680 | 0.1156 |
| **ALB_2** | - | - | - | - | - | 0.0000 | 0.1185 |
| **ALB_3** | - | - | - | - | - | - | 0.0000 |

**Figure S1.** Principal component analysis (PCA) plot illustrating the ordination of samples from Sicilian, Greek and silver firs using the 20 SNPs designed to maximize the species differentiation. Colored dots represent the Sicilian (green), silver (light yellow), and Greek (orange) firs.

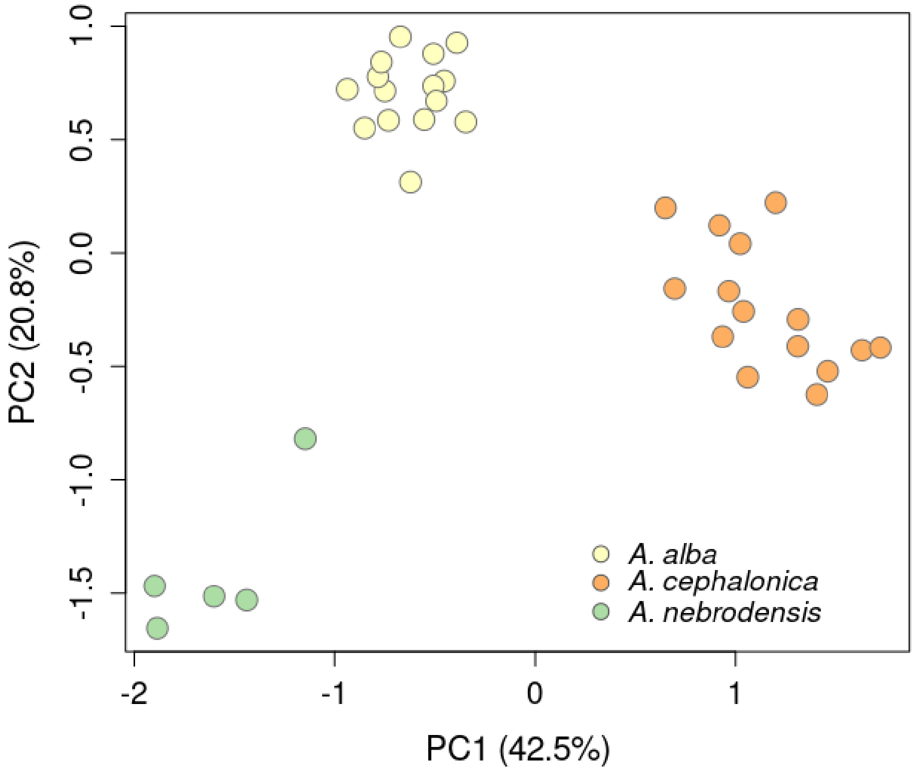

**Figure S2.** Discriminant analysis of principal component (DAPC) and matrix of pairwise co-ancestry among adult trees of the Sicilian fir. The DAPC (a) shows the existence of three genetic clusters, along with the probability of membership for each of the 30 adult trees. The matrix of pairwise co-ancestry among all adult trees (b) is based on Ritland (RIT) estimators, which are represented using relative point size. Colors represent a sequence of RIT values ranging from negative (orange) to positive (blue). Font color (green, red, and blue) corresponds to each of the three previously described genetic clusters. Pairwise comparisons within the same genetic cluster are framed within a box.

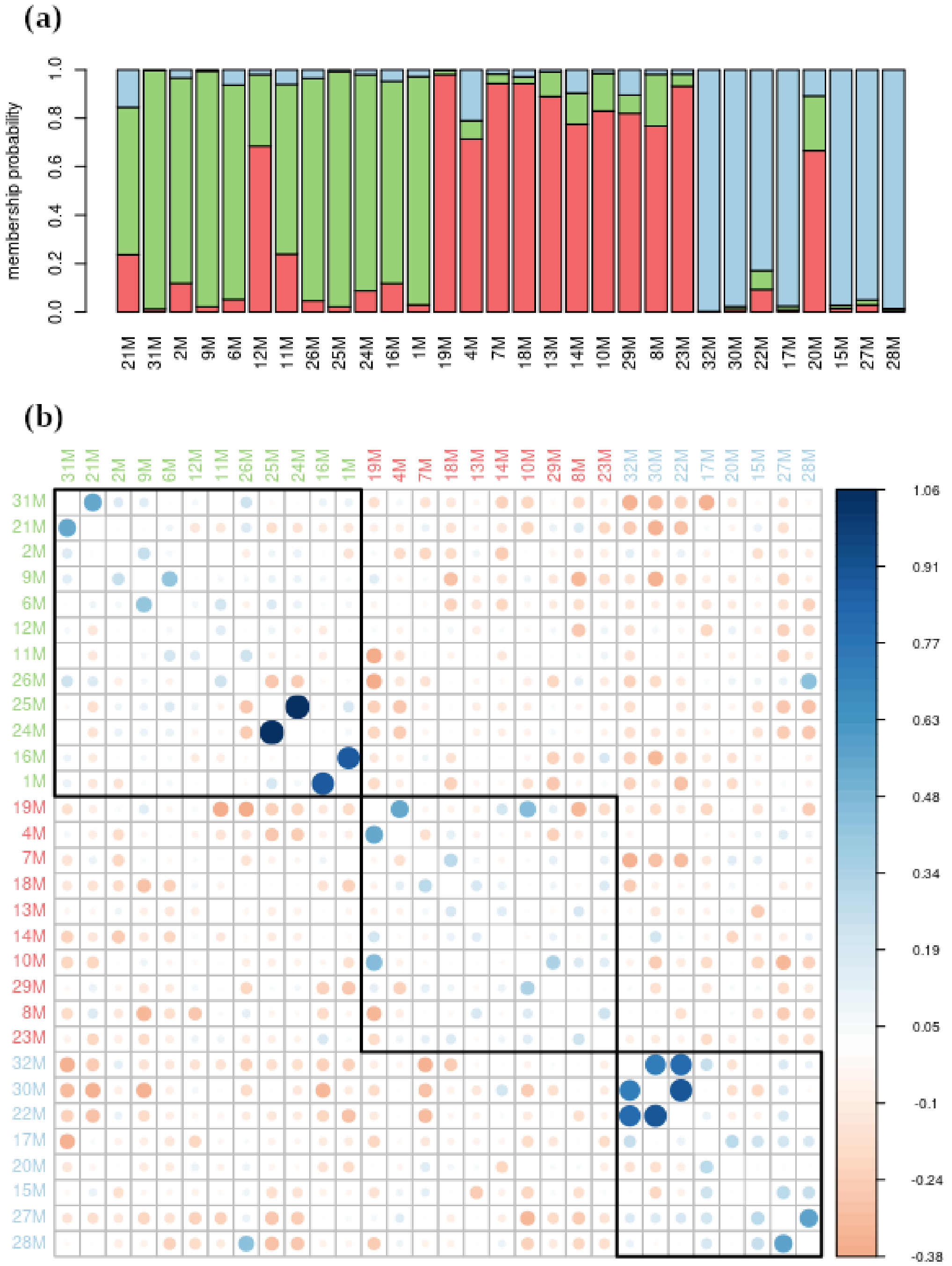
